# Rapid expression of COVID-19 proteins by transient expression in tobacco

**DOI:** 10.1101/2020.12.29.424712

**Authors:** Penelope Lindsay, Amanda Ackerman, Yinan Jian, Oliver Artz, Daniele Rosado, Tara Skopelitis, Munenori Kitagawa, Ullas V. Pedmale, David Jackson

**Author notes:** **Author contributions** P.L., A.A.,Y.J., O.A.,T.S., U.V.P., D.J. designed and performed all experiments and analysis. M.K. performed microscopy, and L.J.T. advised on protein purification, D.J. and U.V.P initiated the project and discussed the results and P.L., U.V.P and D.J. prepared figures and cowrote the manuscript. All authors edited the manuscript.

## Abstract

In 2020 we suffered from a major global pandemic caused by the SARS-CoV-2 coronavirus. Efforts to contain the virus include the development of rapid tests and vaccines, which require a ready supply of viral proteins. Here we report the production of two SARS-CoV-2 proteins by transient transformation of tobacco, leading to high expression within three days, and subsequent purification of the intact proteins. Such efforts may help to develop testing resources to alleviate the major impacts of this global pandemic.

## INTRODUCTION

The COVID-19 pandemic has had a major impact on global health and economic productivity. A major limitation in the management of the pandemic has been limited availability of testing, which can be performed by PCR or by immunological testing. The latter is a better indicator of whether a patient is developing and will maintain immunity to the virus, and requires the production of viral proteins to test for patient antibodies.

Production of heterologous proteins in plants is a well-developed technology that has already been commercialized in several countries. For example, the first FDA approved Ervebo vaccine for the prevention of Ebola virus disease was made in tobacco (Merck). A common approach is to use transient expression in a tobacco species, *Nicotiana benthamiana* (Bally et al., 2015), which is rapid and can be scaled up to commercial scale production (Fischer et al., 2013). In addition, optimized vector systems that boost expression of target proteins have been developed (Norkunas et al., 2018; Sainsbury et al., 2009), leading to the production of up to several grams per kilogram of tobacco tissue. This production requires less specialized infrastructure than other protein production systems, and is easily scalable.

Here we report the production of two SARS-CoV-2 proteins using transient expression in tobacco. We compare different expression systems and codon optimized constructs, and show that high levels of expression can be obtained in just a few days, and that recombinant proteins can be purified using routine epitope tagging. Such approaches can help protein production for basic research as well as applied applications such as development of diagnostics and vaccine production.

## RESULTS

To rapidly express SARS-CoV-2 proteins in plants, we chose *Nicotiana benthamiana* as a convenient expression host. We focused on two surface proteins of the virus, the spike and the nucleocapsid (N) proteins, which are likely to generate immune responses, and therefore most useful for diagnostic testing or vaccine development. For spike protein expression, we based our design on an approach optimized for expression in mammalian cells, lacking the transmembrane domain, and with amino acid substitutions to avoid proteolytic cleavage and to promote assembly of the trimeric spike complex (Wrapp et al., 2020). Since the spike protein is very large, and could pose problems for expression, we also cloned constructs to express the receptor binding domain (RBD) because it is likely to be a primary domain for immune responses. Each construct was generated by amplification from the mammalian codon optimized template, or by synthesis of tobacco codon optimized clones. We used Gateway cloning (Thermo Fisher) to insert the different fragments into a vector along with epitope tags, including mCitrine (mCit), or a modified Flash Tag (2xStrepII-6xHis-3xFlag; Pedmale et al., 2016) and all were expressed under the control of a double cauliflower mosaic virus (CaMV) 35S promoter (2×35S, Figure 1). We also cloned fragments containing a 8xHis-2xStrepII tag into the pEAQ.HT DEST1 vector, which is designed to maximize expression in *N. benthamiana* (Sainsbury et al., 2009). After sequence verification, the constructs were introduced into *Agrobacterium,* and cultures used to infiltrate tobacco plants, either by syringe infiltration or by submerging whole plants in an *Agrobacterium* suspension in a vacuum chamber (Stephenson et al., 2018). Following infiltration, plants were sampled at different time points to find the optimal expression of the tagged proteins, either by confocal microscopy for the mCit tagged clones (Figure 2), or by western blotting (Figure 3). For mCit tagged spike constructs, we observed the “full length” protein at the cell periphery, as well as in a perinuclear and reticulate pattern, reminiscent of endoplasmic reticulum, as expected. However, it was not clear if the protein was secreted, because it remained associated with the plasma membrane following plasmolysis (Figure 2C). The RBD-mCit fusion protein appeared to localize cytoplasmically, in as yet unidentified mobile puncta (Figure 2E, Supp movie). We used western blotting to confirm that the proteins were intact and of the expected size, and also to find the optimal expression period. Expression was highest at three days after infiltration when spike was expressed in pB7m34GW, and at five days when expressed in pEAQ.HT. In summary, full length and RBD domain spike proteins accumulated to high levels in tobacco, and were correctly localized.

**Figure 1.**
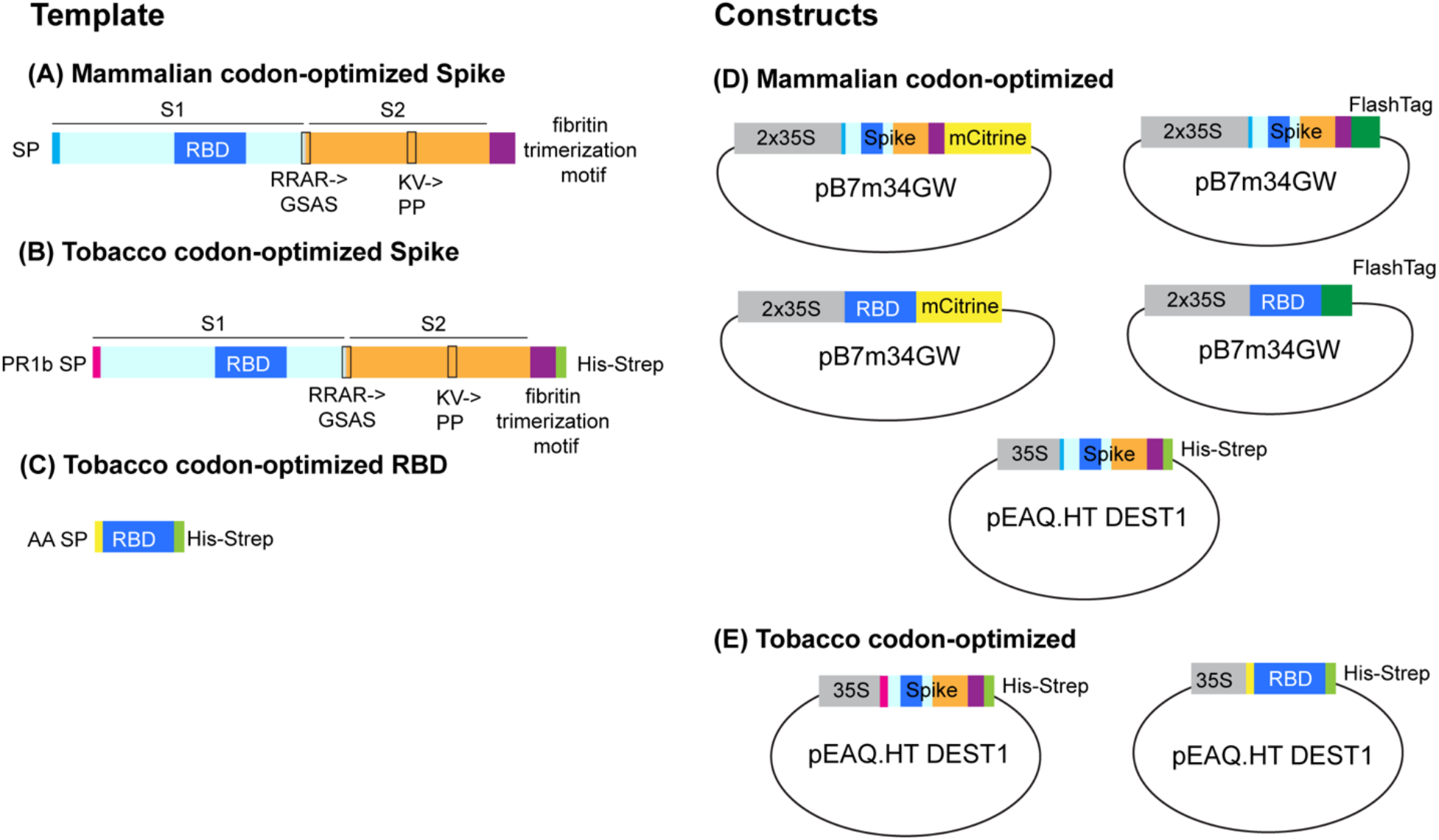
Spike constructs generated in this study. Mammalian codon-optimized (A), or tobacco codon-optimized (B - C) spike ectodomain coding sequences were used to clone constructs. All constructs lacked the C-terminal transmembrane domain and have amino acid substitutions that abolish the proteolytic cleavage domain between the S1 and S2 subunits (RRAR>GSAS). A c-terminal fibritin trimerization motif (purple) promotes trimerization in the absence of the transmembrane domain. The tobacco codon-optimized full-length spike has a pathogenesis-related 1b (Pr1b) signal peptide (pink) in place of the native spike signal peptide (SP), and the RBD domain has an alpha amylase (AA) signal peptide (yellow). Mammalian codon-optimized spike coding sequences were cloned into pB7m34GW destination vector with a 2×35S promoter and either a C-terminal mCitrine or FlashTag (containing a Strep-His-Flag epitope, D). The mammalian-optimized full-length spike CDS with a C-terminal 8xHis-2xStrepII tag was also cloned into pEAQ.HT DEST1, which contains P19 and cowpea mosaic viral UTRs to maximize expression in tobacco. Tobacco codon-optimized sequences were cloned into pEAQ.HT DEST1 (E), with a C-terminal 8xHis-StrepII tag, to maximize expression levels.

**Figure 2.**
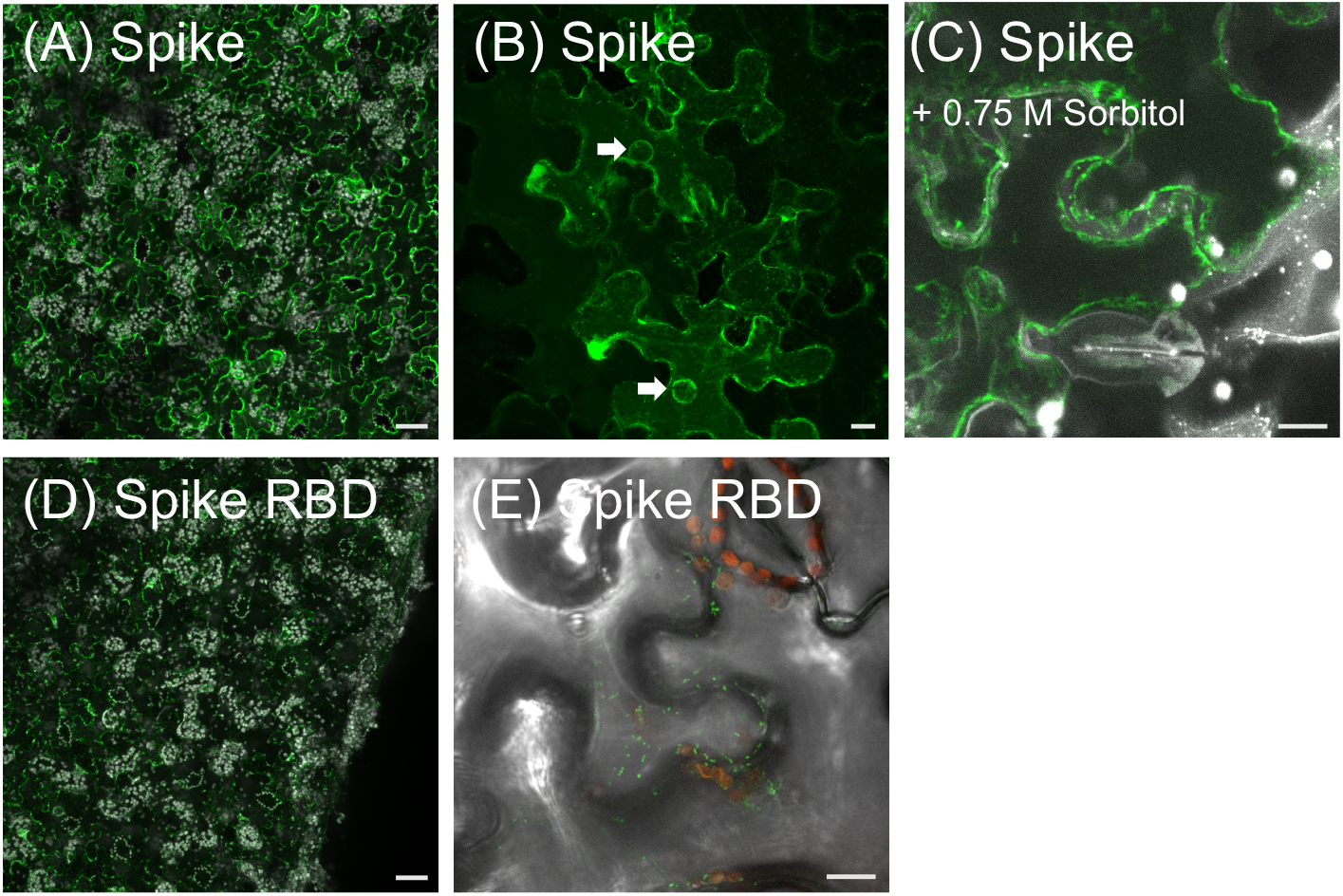
Visualization of mCitrine-tagged SARS-CoV-2 spike proteins in tobacco. *N. benthamiana* leaves were syringe infiltrated with *Agrobacterium* containing spike ectodomain or RBD domains and imaged using confocal microscopy. (A) – (C) Green, Spike-mCitrine fusion protein signal, associated with perinuclear ring (arrows), likely ER, and plasma membrane, that separates from the cell wall following plasmolysis. (D) – (E) mCitrine-tagged RBD proteins localized to cytoplasmic puncta, enlarged in (E). In (A), (D) grey = chloroplast autofluorescence, in (C) grey = propidium iodide counterstain, (E) red = chloroplast autofluorescence, grey = DIC. In (A), (D), scale bars are 20 microns. (B), (C), (E), scale bars are 10 microns.

**Figure 3.**
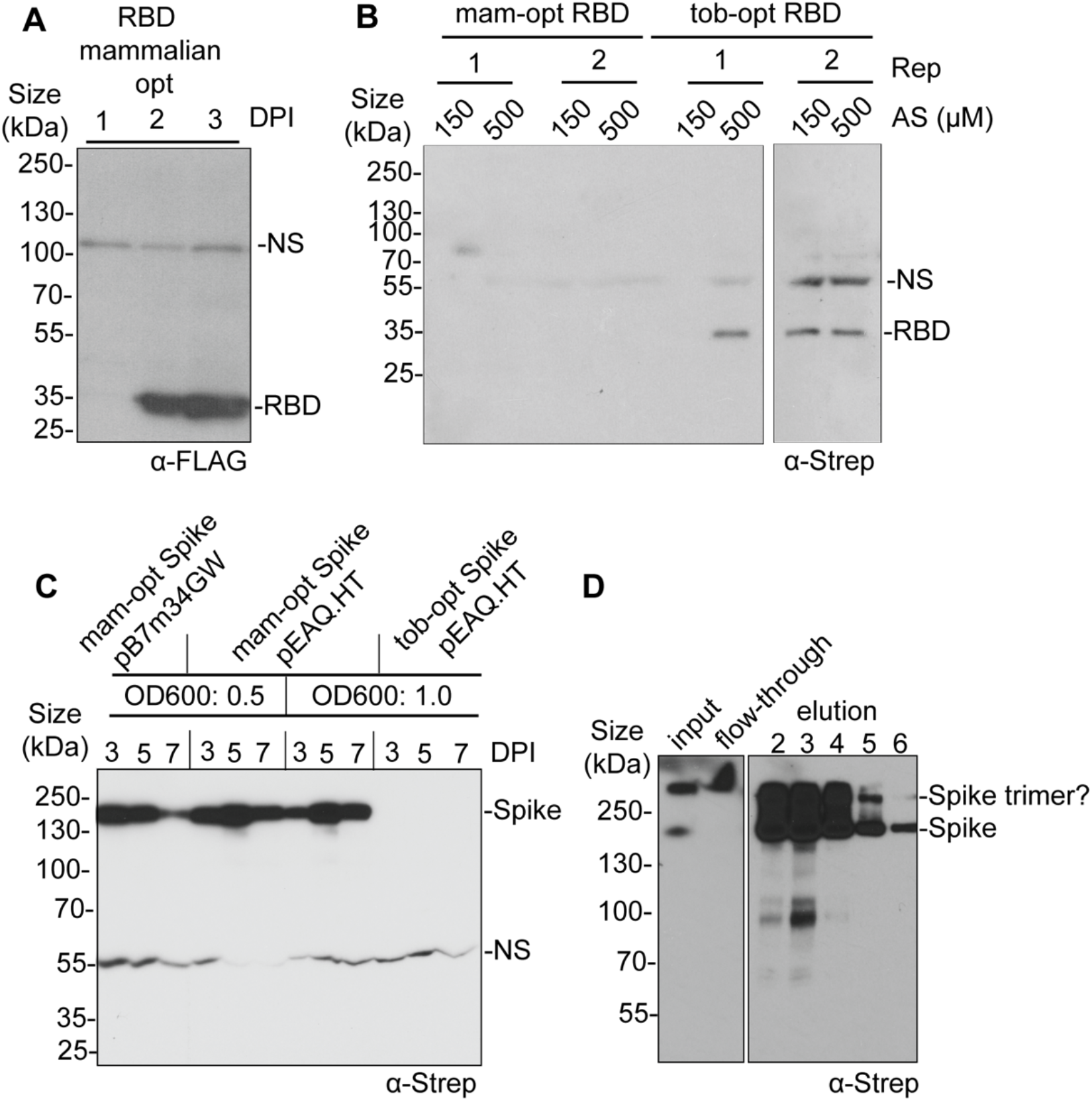
Expression and purification of spike proteins in tobacco. (A) Mammalian codon-optimized RBD proteins were detected using anti-FLAG antibody and were higher at three days post infiltration (DPI) and guide optimal expression conditions. (B) Expression levels of *N. benthamiana* codon-optimized (tob-opt) RBD in the pEAQ.HT expression vector were higher than the mammalian codon-optimized version (mam-opt) in pB7m34GW. Proteins were detected using the anti-Strep antibody, which is less sensitive than anti-FLAG. (C) Expression levels of mammalian-optimized spike ectodomain were comparable between pEAQ.HT and pB7m34GW. The tobacco-codon optimized spike ectodomain in pEAQ.HT DEST1 was expressed lower than the mammalian-optimized spike. Proteins were detected using anti-Strep antibody. NS, nonspecific band, DPI, days post infiltration, AS, acetosyringone concentration. (D), For affinity purification, total protein was extracted from *N. benthamiana* leaves vacuum-infiltrated with *Agrobacterium* containing the mammalian-optimized spike-His-Strep, and spike protein was purified using Strep-Tactin XT resin. Dashes indicate putative trimer and monomeric bands. Spike protein was detected via western blot using the anti-Strep antibody.

We next asked if codon optimization could enhance production of spike protein. The fulllength and RBD coding sequences were optimized to match tobacco codon usage, resulting in a score of 0.78, compared to 0.64 for the mammalian optimized clone. We also changed the signal peptide from the original viral one, to ones that are efficiently processed in tobacco; from alpha amylase for the RBD domain or Pathogenesis-related protein 1b (PR1b) for full-length spike protein (Streatfield et al., 2001; Tuboly et al., 2000). The tobacco codon optimized clones were next inserted into the same expression vector, and infiltrated into tobacco plants. Protein expression using the tobacco-codon optimized RBD clone was higher than that of the mammalian optimized clone, as expected (Figure 3B). To our surprise, however, expression of the full-length tobacco optimized spike protein was lower than that of the mammalian optimized version (Figure 3C), so we proceeded to use the latter for affinity purification. We next vacuum infiltrated whole tobacco plants to collect several grams of expressing tissue, and used Strep-Tactin XT resin to purify spike proteins, according to the manufacturer’s instructions. Despite using tens of g of tissue, purification was inefficient, leading to ~ 100 μg of purified protein. In several purification attempts, we recovered an average of ~7.5 μg of spike protein /g of tissue. Although purification was inefficient, spike protein appeared to be present as a trimer (Figure 3D), suggesting that it folded properly. We observed trimer formation even under denaturing conditions because the fibritin trimerization motif is resistant to denaturing agents such as high temperature and B-mercaptoethanol (Yang et al., 2002).

We used a similar approach for expression of N protein. N-mCitrine was readily observed in tobacco leaf epidermal cells 1-day post infiltration (Figure 4A) by fluorescence wide-field or confocal microscopy, and localized in both the cytoplasm and the nucleus. The nuclear N-mCit protein was often found in puncta (Figure 4A). *N. benthamiana* also expressed high levels of N-Flash protein, but attempts to purify it using its strepII-tag with the Strep-Tactin resin failed. Similarly, efforts to purify it using its 6xHis-tag with cobalt IMAC beads under a variety of conditions were unsuccessful. It is likely that the epitope tag in N-Flash was not accessible in our capture methods, and therefore we used a C terminally tagged Flash-N construct for subsequent purifications. Flash-N (Figure 4C) expressed readily, similar to N-Flash and N-mCitrine, and high levels of expression were detected (Figure 4C). We employed the strepII-tag on Flash-N and Strep-tactin resin to purify it from total protein lysates prepared from leaves 3-days post infiltration. We observed improved binding and subsequent purification of Flash-N protein (Figure 4D), indicating that the position of the tag is important for purification from *in planta* systems.

**Figure 4.**
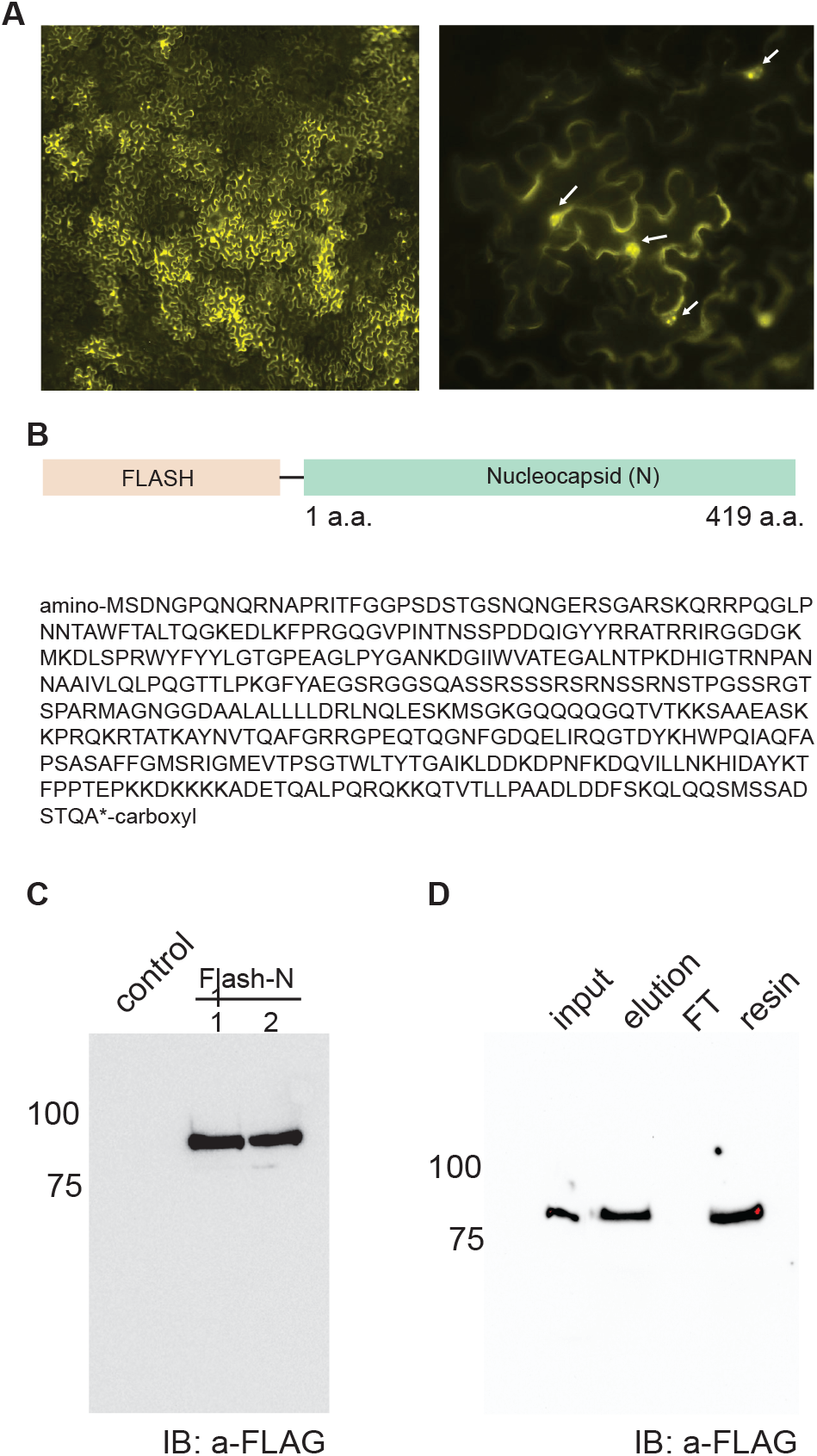
Expression and detection of N protein. a) Expression of N-mCitrine protein in *N. benthamiana* leafepidermal cells 1-day post infiltration. Arrowsi ndicatethenucleus.b) schematic diagram of Flash-N pratein alongwithth) amino acid sequence. c) Immunoblot analysis reveals expression of Flash-N in two *independent N*. benthamiana leaves from different plants (1 and 2) after 1-day post infiltration. Immunoblot was performed on total proteins using anti-Flag antibodies. d) Immunoblot analysis using anti-Flag antibody on the purification of Flash-N protein using Strep-Tactin XT resin. FT indicates flow-through proteins that did not bind to the resin. Elution refers to the proteins eluted from the resin using biotin and the resin lane indicates proteins bound to it.

In summary, we show that we can successfully express SARS-Cov-2 proteins in tobacco. Other recent reports (Diego-Martin et al., 2020; Makatsa et al., 2020; Margolin et al., 2020; Rattanapisit et al., 2020) indicate similar results. The rapid and high level of expression makes this an attractive system for larger scale production for research, or development of tests or of vaccines.

## DISCUSSION

In this study, we used *N. benthamiana* plants as a heterologous system to produce recombinant SARS-CoV-2 proteins. The rapid expression, ability to produce large amounts of protein, and easily scalable system make this an attractive method to generate SARS-CoV-2 proteins for diagnostic tests and as potential vaccine candidates (Capell et al., 2020).

We successfully purified recombinant spike and Nucleocapsid SARS-CoV-2 proteins from *N. benthamiana* at a concentration comparable to other studies (Diego-Martin et al., 2020; Makatsa et al., 2020; Margolin et al., 2020; Rattanapisit et al., 2020). The spike ectodomain was purified in both monomeric and trimeric forms, suggesting that it was folding properly. Additionally, we confirmed that the spike ectodomain localized to the ER, where protein glycosylation occurs (Strasser, 2016). The spike protein is heavily glycosylated, and future work should examine whether these modifications occur correctly in plants (Watanabe et al., 2020). Together, our data support the idea that plants are a suitable eukaryotic system for recombinant production of SARS-CoV-2 proteins. Indeed, other studies have used plant-produced SARS-CoV2 antigens in ELISA assays to detect human antibodies, as well as monoclonal antibodies produced *in planta,* demonstrating their efficacy in diagnostics (Diego-Martin et al., 2020; Makatsa et al., 2020; Rattanapisit et al., 2020).

We were surprised that a tobacco codon optimized spike construct expressed at lower levels than the mammalian codon optimized version, but several other reports expressing mammalian proteins in tobacco have yielded similar results (Maclean et al., 2007; Margolin et al., 2020; Mortimer et al., 2012). In contrast, however, the tobacco codon-optimized RBD construct produced more protein than the mammalian codon-optimized version. The two tobacco codon-optimized spike constructs were synthesized with different signal peptides, which could explain the difference in protein accumulation. We also tested two vector backbones, pEAQ.HT, which contains a single 35S promoter, and pB7m34GW with 2×35S promoter, and found that spike ectodomain expression levels were similar. This was surprising, since pEAQ.HT contains the 5’ and 3’ UTRs of cowpea mosaic virus to boost expression levels (Sainsbury et al., 2009). Despite the fact that expression levels were similar, pEAQ.HT has the P19 gene silencing suppressor incorporated into it, making it simpler to work with for large scale infiltrations.

While our yield of SARS-CoV-2 proteins from tobacco was modest compared to other expression systems, plants still offer several advantages as a heterologous system for protein production. Producing recombinant proteins in plants can easily and affordably be scaled up compared to other systems, offering decentralized capabilities in times of rapid demand for diagnostics (Diego-Martin et al., 2020). Although more research is needed to fully understand the dynamics of COVID19 immunity, recent studies indicate that protective immunity may only last over a period of months (Poland et al., 2020). This suggests we may need repeated vaccinations to offer protective immunity to a global population, necessitating affordable vaccine candidates. Future work should also explore the potential of plant-based oral vaccines to combat SARS-CoV-2, as these are a cost-effective alternative suitable for repeated dosages (Chan and Daniell, 2015).

## METHODS

### Cloning

We used Gateway cloning (Thermo Fisher) to generate spike and Nucleocapisd constructs for expression in *N. benthamiana*. Mammalian codon-optimized entry clones were generated by PCR amplification of spike ectodomain vector as a template (Wrapp et al., 2020) using Phanta High Fidelity polymerase (Vazyme). This mammalian codon-optimized sequence contained the amino acid substitutions described in Wrapp et al., 2020. Primers PL58 (GGGGACAAGTTTGTACAAAAAAGCAGGCTTCatgttcgtgttcctggtgct) and PL65 (GGGGACCACTTTGTACAAGAAAGCTGGGTCggatctgcccaggaatgtgc) were used to amplify the spike ectodomain and PL74 (GGGGACAAGTTTGTACAAAAAAGCAGGCTTCATGccaaacattaccaacctctgcc) and PL75 (GGGGACCACTTTGTACAAGAAAGCTGGGTCgggggcgtggagtaactc) for the RBD domain. *N. benthamiana* codon-optimized clones were codon-optimized using Genscript and synthesized using Gene Universal. PCR amplified products containing attB recombination sites were gel purified and recombined into pDONR 1-2 or pDONR207 using BP clonase. Entry clones were verified by sequencing the insert and restriction digest.

For multisite Gateway cloning, pDONR-P4P1R containing 2xCaMV 35S promoter, pDONR-221 with spike, and either pDONR-P2RP3 mCitrine or FlashTag entry clones were recombined into the pB7m34GW (Pedmale et al. 2016) binary destination vector using LR clonase (Thermo Fisher). Destination clones were verified by restriction digest and sequencing of insertion borders. *N. benthamiana* or mammalian codon optimized sequences containing the spike ectodomain or RBD with C-terminal 8xHis-Twin Strep tags were cloned into pEAQ.HT DEST1 using LR clonase (Sainsbury et al., 2009). Destination clones were transformed into the *Agrobacterium tumefaciens* strain GV3101.

Full-length N protein was first amplified from the reverse transcribed product of the above synthetic viral mRNA using oligos AA17 (5’-GGGGACAGCTTTCTTGTACAAAGTGGccATGtctgataatggac-3’) and AA18 (5’-GGGGACAACTTTGTATAATAAAGTTGcTCAggcctgagttgagtcag-3’) containg attB2 and attB3 Gateway recombination sites. The PCR amplicon was then cloned in to pDONR-P2RP3 using BP II clonase (Thermo Fisher). The resulting clone was then recombined with pB7m34GW destination vector using LR II clonase (Thermo Fisher) along with 2xCaMv35S promoter in pDONR-P4P1R and Flash-tag in pDONR-221 to obtain *2×35Spro::Flash-N.* Similarly, *2×35Spro::N-mCitrine and 2×35Spro::N-Flash* constructs were generated.

### *N. benthamiana* propagation and infiltrations

Four-to five-week old *N. benthamiana* plants grown in long-day growth conditions were infiltrated with *Agrobacterium tumefaciens* suspended to an optical density of 0.5 – 1.0 in infiltration medium (10 mM MES, pH 5.6, 10 mM MgCl2) containing 150 μM acetosyringone using either syringe or vacuum infiltration at 2 days post infiltration, unless otherwise indicated. For vacuum infiltrations, whole plants were submerged in agrobacterium suspensions in a vacuum chamber, and a vacuum pressure of approx. 30 in. Hg was applied for 3 minutes, until all leaves were visibly infiltrated with the bacterial solution.

The pB7m34GW constructs were co-infiltrated with p19 expressing constructs to suppress gene silencing in *N. benthamiana.*

### Confocal microscopy

*N. benthamiana* leaves transiently expressing SARS-CoV-2 proteins were imaged using a Zeiss LSM 900 confocal microscope. The objectives used for imaging were as follows: EC Plan-Neofluar 10x/0.30 NA M27, Plan-Apochromat 20x/0.8 NA M27, LD LCI Plan-Apochromat 40x/1.2 NA Imm Korr DIC M27, C Plan-Apochromat 63x/1.4 NA.

For plasmolysis experiments, leaves were incubated in 0.75 M sorbitol for 15 minutes, and counterstained with propidium iodide to visualize the plasma membrane. mCitrine was imaged using 488 nm excitation, and the emission wavelength collected was from 410 – 546 nm. Chlorophyll autofluorescence was imaged using 640 nm excitation, and emission wavelength collected was from 650 – 700 nm. Propidium iodide was excited at 561 nm, and emission wavelength collected was from 598 – 617 nm.

### Protein extraction, affinity purification, and immunoblotting

To assess protein levels, two 0.75 mm leaf punches were ground and boiled for 10 min in 75 μL of 2X SDS loading buffer containing 5 % B-mercaptoethanol. 20 μL of the crude extract was loaded onto a 1 mm thick 8% SDS-PAGE gel, run for 2 hours at 80 V, and then transferred onto PVDF membrane (Immobilon) for 30 V overnight at 4 ° C. Blots were dehydrated with a 10 second incubation in methanol, followed by drying for 15 min at room temperature. They were next incubated in primary antibody solution for 2 hours at room temperature (1:1000 anti-Strep, IBA Life Sciences, 1:10,000 anti-FLAG M2 monoclonal antibody, Sigma in 2% milk Tris – buffered saline, 0.1% Tween (TBST)), followed by three ten minutes washes in TBST. Then, blots were incubated for 1 hour in the secondary antibody solution (ECL Anti-mouse conjugated to horseradish peroxidase, 1:10,000, GE in 2% milk), followed by three ten-minute washes in TBST. Immunoblots were developed using Amersham Prime ECL Western blotting reagent (GE) and imaged using autoradiography film.

For spike protein affinity purification, frozen *N. benthamiana* tissue was ground using either a mortar and pestle or Nutriblend bullet blender (2 min blend) in three volumes of extraction buffer per gram tissue (50 mM Tris pH 8, 150 mM NaCl, 1% Triton X, protease inhibitor cocktail). Samples were centrifuged for 5 minutes at 1000 g, and the supernatant was passed through Miracloth to remove debris. The extract was centrifuged at 20,000 g for 15 min, and the supernatant loaded onto a column containing Strep-Tactin XT resin (IBA Life Sciences) and protein was purified according to the manufacturer’s protocol.

<u>Purification and detection of N protein</u>. 3 days post infiltration, *N. benthamina* leaves were flash frozen in liquid N2 and then ground to a fine powder using a mortar and pestle and resuspended 1:4 ratio in RIPA buffer (25 mM Tris-HCl, pH 7.6, 150 mM NaCl, 1% NP-40, 1% sodium deoxycholate, 0.1% SDS) supplemented with 1’ protease inhibitor cocktail (Millipore Sigma) and 100 μg/avidin. The lysate was gently rotated for 5 min at 4°C and then sonicated at 45% intensity, with 0.5 second on and off pulses for a total of 30 seconds. The lysate was then filtered through miracloth and centrifuged at 20,000×g for 15 minutes. Pre-washed Strep-Tactin XT resin (IBA Life Sciences) was then added to the cleared lysate and gently rotated at 4°C for 1 hr. The resin was then washed 3 times with RIPA buffer and eluted with 1’ Strep-Tactin elution buffer with biotin (IBA Life Sciences). The input, flow-through and eluted protein samples were then boiled with 2× LDS sample buffer for 5 minutes before performing Bis-Tris polyacrylamide gel electrophoresis (PAGE).

After SDS-PAGE, the proteins were immobilized on to a reinforced nitrocellulose membrane (Millipore Sigma) with 1× NuPage transfer buffer (Thermo Fisher) containing 10% methanol. The membrane was blocked with 5% fat-free milk prepared in TBST (20 mM Tris-HCl, pH 7.6, 137 mM NaCl, and 0.05% Tween 20) for 10 minutes and then incubated for 1 hr with anti-Flag-HRP (Sigma) at 1:5000 dilution in 1% milk in TBST with gentle shaking at room temperature. The membrane was then washed three times for 10 minutes each with TBST and

Flag-specific bands were detected by chemiluminescence using appropriate substrate (SuperSignal West Dura; Thermo Fisher).

## Acknowledgments

The authors would like to acknowledge funding from CSHL, and from grants for the National Science Foundation (IOS-1930101and IOS-1755141 to D.J). U.V.P. acknowledges NIH grants GM125003 and GM125003-03S1. We thank Tim Mulligan and Kyle Schlecht for plant care, Leemor Joshua-Tor for the mammalian-optimized spike ectodomain vector and Elad Elkayam for assistance with large-scale affinity purifications.

## References

Bally, J., Nakasugi, K., Jia, F., Jung, H., Ho, S.Y.W., Wong, M., Paul, C.M., Naim, F., Wood, C.C., Crowhurst, R.N., et al. (2015). The extremophile Nicotiana benthamiana has traded viral defence for early vigour. Nat. Plants 1, 15165.

Capell, T., Twyman, R.M., Armario-Najera, V., Ma, J.K.-C., Schillberg, S., and Christou, P. (2020). Potential Applications of Plant Biotechnology against SARS-CoV-2. Trends Plant Sci. 25, 635–643.

Chan, H.-T., and Daniell, H. (2015). Plant-made oral vaccines against human infectious diseases-Are we there yet? Plant Biotechnol. J. 13, 1056–1070.

Diego-Martin, B., González, B., Vazquez-Vilar, M., Selma, S., Mateos-Fernández, R., Gianoglio, S., Fernández-del-Carmen, A., and Orzáez, D. (2020). Pilot production of SARS-CoV-2 related proteins in plants: a proof of concept for rapid repurposing of indoors farms into biomanufacturing facilities. BioRxiv 2020.10.13.331306.

Fischer, R., Schillberg, S., F. Buyel, J., and M. Twyman, R. (2013). Commercial Aspects of Pharmaceutical Protein Production in Plants. Curr. Pharm. Des. 19, 5471–5477.

Maclean, J., Koekemoer, M., Olivier, A.J., Stewart, D., Hitzeroth, I.I., Rademacher, T., Fischer, R., Williamson, A.-L., and Rybicki, E.P. (2007). Optimization of human papillomavirus type 16 (HPV-16) L1 expression in plants: comparison of the suitability of different HPV-16 L1 gene variants and different cell-compartment localization (Microbiology Society,).

Makatsa, M.S., Tincho, M.B., Wendoh, J.M., Ismail, S.D., Nesamari, R., Pera, F., de Beer, S., David, A., Jugwanth, S., Gededzha, M.P., et al. (2020). SARS-CoV-2 antigens expressed in plants detect antibody responses in COVID-19 patients. MedRxiv 2020.08.04.20167940.

Margolin, E., Verbeek, M., Meyers, A., Chapman, R., Williamson, A.-L., and Rybicki, E.P. (2020). Calreticulin co-expression supports high level production of a recombinant SARS-CoV-2 spike mimetic in *Nicotiana benthamiana*. BioRxiv 2020.06.14.150458.

Mortimer, E., Maclean, J.M., Mbewana, S., Buys, A., Williamson, A.-L., Hitzeroth, I.I., and Rybicki, E.P. (2012). Setting up a platform for plant-based influenza virus vaccine production in South Africa. BMC Biotechnol. 12, 14.

Norkunas, K., Harding, R., Dale, J., and Dugdale, B. (2018). Improving agroinfiltration-based transient gene expression in Nicotiana benthamiana. Plant Methods 14, 71.

Pedmale, U.V., Huang, S.C., Zander, M., Cole, B.J., Hetzel, J., Ljung, K., Reis, P.A.B., Sridevi, P., Nito, K., Nery, J.R., et al. (2016). Cryptochromes Interact Directly with PIFs to Control Plant Growth in Limiting Blue Light. Cell 164, 233–245.

Poland, G.A., Ovsyannikova, I.G., and Kennedy, R.B. (2020). SARS-CoV-2 immunity: review and applications to phase 3 vaccine candidates. The Lancet 396, 1595–1606.

Rattanapisit, K., Shanmugaraj, B., Manopwisedjaroen, S., Purwono, P.B., Siriwattananon, K., Khorattanakulchai, N., Hanittinan, O., Boonyayothin, W., Thitithanyanont, A., Smith, D.R., et al. (2020). Rapid production of SARS-CoV-2 receptor binding domain (RBD) and spike specific monoclonal antibody CR3022 in Nicotiana benthamiana. Sci. Rep. 10, 17698.

Sainsbury, F., Thuenemann, E.C., and Lomonossoff, G.P. (2009). pEAQ: versatile expression vectors for easy and quick transient expression of heterologous proteins in plants. Plant Biotechnol. J. 7, 682–693.

Stephenson, M.J., Reed, J., Brouwer, B., and Osbourn, A. (2018). Transient Expression in Nicotiana Benthamiana Leaves for Triterpene Production at a Preparative Scale. J. Vis. Exp. e58169.

Strasser, R. (2016). Plant protein glycosylation. Glycobiology 26, 926–939.

Streatfield, S.J., Jilka, J.M., Hood, E.E., Turner, D.D., Bailey, M.R., Mayor, J.M., Woodard, S.L., Beifuss, K.K., Horn, M.E., Delaney, D.E., et al. (2001). Plant-based vaccines: unique advantages. Vaccine 19, 2742–2748.

Tuboly, T., Yu, W., Bailey, A., Degrandis, S., Du, S., Erickson, L., and Nagy, E. (2000). Immunogenicity of porcine transmissible gastroenteritis virus spike protein expressed in plants. Vaccine 18, 2023–2028.

Watanabe, Y., Allen, J.D., Wrapp, D., McLellan, J.S., and Crispin, M. (2020). Site-specific glycan analysis of the SARS-CoV-2 spike. Science 369, 330.

Wrapp, D., Wang, N., Corbett, K.S., Goldsmith, J.A., Hsieh, C.-L., Abiona, O., Graham, B.S., and McLellan, J.S. (2020). Cryo-EM structure of the 2019-nCoV spike in the prefusion conformation. Science 367, 1260.

Yang, X., Lee, J., Mahony, E.M., Kwong, P.D., Wyatt, R., and Sodroski, J. (2002). Highly Stable Trimers Formed by Human Immunodeficiency Virus Type 1 Envelope Glycoproteins Fused with the Trimeric Motif of T4 Bacteriophage Fibritin. J. Virol. 76, 4634.

